# Insulinase like protease 1 contributes to macrogamont formation in *Cryptosporidium parvum*

**DOI:** 10.1101/2020.12.03.411165

**Authors:** Rui Xu, Yaoyu Feng, Lihua Xiao, L. David Sibley

## Abstract

The apicomplexan parasite *Cryptosporidium parvum* contains an expanded family of 22 insulinase like proteases (INS), a feature that contrasts with their otherwise streamlined genome. Here we examined the function of INS1, which is most similar to the human insulinase protease that cleaves a variety of small peptide substrates. INS1 is a M16A clan member and contains a signal peptide, an N-terminal domain with the HxxEH active site, followed by three inactive domains. Unlike previously studied *C. parvum* INS proteins that are expressed in sporozoites and during merogony, INS1 was expressed exclusively in macrogamonts, where it was localized in small cytoplasmic vesicles. Although INS1 did not colocalize with the oocyst wall protein recognized by the antibody OW50, immune-electron microscopy indicated that INS1 resides in small vesicles in the secretory system. Notably, these small INS1 positive vesicles often subtend large vacuoles resembling wall forming bodies, which contain precursors for oocyst wall formation. Genetic deletion of INS1, or replacement with an active site mutant, resulted in lower formation of macrogamonts *in vitro* and reduced oocyst shedding *in vivo*. Our findings reveal that INS1 functions in formation or maturation of macrogamonts and that its loss results in attenuated virulence in immunocompromised mice.

**Importance:** Cryptosporidiosis is a debilitating diarrheal disease in young children in developing countries. Absence of effective treatments or vaccines makes this infection very difficult to manage in susceptible populations. Although the oral dose of oocysts needed to cause infection is low, infected individuals shed very high numbers of oocysts, hence readily contaminating the environment. Our studies demonstrate that the protease INS1 is important for formation of female sexual stages and that in its absence, parasites produce fewer oocysts and are attenuated in immunocompromised mice. These findings suggest that mutants lacking INS1, or related proteases, may be useful for producing attenuated vaccines to induce immunity without causing disease.

---

*Cryptosporidium* spp. are apicomplexan parasites that cause diarrheal disease in humans and animals. Human infection is primarily caused by two species, *C. parvum* that also infects agricultural ruminant animals and is zoonotic, and *C. hominis* that is spread human-to-human (1). Cryptosporidiosis was recognized as one of the top three causes of severe diarrhea in children younger than two years of age in developing countries as reported by the Global Enteric Multi-Center Study (2). Infection in early life is also associated with lasting defects in development even after infections subside (3). Nitazoxanide is the only FDA approved drug for the treatment cryptosporidiosis. However, it has a limited effect in immunocompromised individuals and is not approved for use in children under the age of two (4). There are currently no effective vaccines for *C. parvum* and fundamental studies on parasite biology and host-pathogen interactions are needed to identify potential therapeutic and vaccine targets.

The entire life cycle of *Cryptosporidium* occurs in a single host leading to efficient fecal-oral transmission (5). Following ingestion of oocysts, sporozoites emerge and invade intestinal epithelial cells where they develop in a unique vacuole formed at the apex of the host cell (6). The parasite initially grows as a trophozoite before undergoing multiple rounds of asexual replication during merogony (7, 8). The parasite then differentiates to the sexual phase and develops as macrogamonts or microgamonts that begin appearing after 44-48 hours post-infection (hpi) (7, 8). However, in transformed cell lines grown *in vitro*, the infection does not progress and parasite numbers gradually decline (9). Comparison of *in vitro* cultures in adenocarcinoma cells lines to the developmental process that occurs in the intestine of mice revealed that the block to complete development *in vitro* is due to lack of fertilization, despite the fact that both gametocyte forms develop normally (10).

Biological investigations of *Cryptosporidium* have been hampered by limitations in experimental platforms for *in vitro* growth. Despite this limitation, *in vitro* propagation systems have been used to generate antibodies that identify different stages of *C. parvum* (11), leading to a better understanding the lifecycle (12). Recent developments have also provided systems that allow complete development of infectious oocysts in stem-cell derived cultures *in vitro* (13, 14). Finally. advances in CRISPR/Cas9 technology have allowed genetic modification in *C. parvum* to tag genes for localization and disrupt them to study function (15). Despite these advances, we lack an understanding of the function of most genes in *Cryptosporidium*, many of which have no orthologues outside the genus, or which are specific for the phylum Apicomplexa (16, 17).

The genome of *C. parvum* is highly streamlined with short intergenic regions, limited introns, and loss of many metabolic pathways (18). Greater than 98% of genes in *C. parvum* are present as a single copy, and only there are a limited number of multigene families that include insulinase-like proteases (INS), MEDLE proteins, and mucin-type glycoproteins (19). INS proteins belong to the M16 family of metallopeptidases that play diverse roles in and they can be found in the cytosol, organelles, and even the cell surface (20). M16 metalloproteases typically bind zinc as part of their active site (HXXEH) and they cleave short polypeptides, the size of which is constrained by a conserved small barrel fold that forms the catalytic chamber (21). The apicomplexan parasite *Toxoplasma gondii* contains ~ 50 metalloproteases, including 11 members of the M16 clan (22), several of which are found in secretory organelles implicated in host cell interactions (23, 24). In *Plasmodium falciparum* the M16 metalloprotease falcilysin participates in both hemoglobin degradation and in processing of transit peptides for apicoplast proteins (25, 26). Similarly, other M16 family members are known for their roles in processing transit peptides for mitochondria and chloroplasts (20).

The *C. parvum* genome contains 22 members of the M16 family of metallopeptidases. Ten of these contain the active site “HXXEH” indicating they might act as enzymes in parasite, although none of their substrates have been defined. Previous studies have shown that antibodies against INS20-19 (although originally annotated as two genes, later assemblies collapsed these into a single gene), INS15, or INS5 inhibit parasite invasion of host cells *in vitro*, suggesting they are involved processing substrates important for host cell recognition or entry (27–29). In the present study, we focused on INS1, which is present among all *Cryptosporidium* spp. INS1 is a classic M16A family member with one active functional motif “HXXEH” followed by three inactive domains. We demonstrate that INS1 is expressed exclusively in macrogamonts of *C. parvum* where is localizes to small transport vesicles. Deletion of INS1, or replacement with an inactive site mutant, reduced formation of macrogamonts *in vitro* and attenuated infection *in vivo*, demonstrating that it is required for optimal oocyst formation.

## Results

### Epitope tagging of *C. parvum* INS1

We compared the sequences of the 22 members of the M16 metalloprotease family present in *C. parvum* using a neighbor joining phylogenetic analysis. For comparison, we also included the human insulinase gene IDE (P14735). Phylogenetic analysis indicated that INS1, encoded by the *cgd1_1680* gene, is most similar to IDE (Figure 1A). INS1 contains a signal peptide and has four domains, one active domain containing zinc-binding motif “HLIEH” and three inactive domains consistent with it belonging to the M16A clan (Figure 1B). To investigate the cellular localization of INS1, we used CRISPR/Cas9 genome editing to tag INS1 with a triple hemagglutinin (3HA) epitope tag at the C-terminus. The tagging construct also contained a selection cassette consisting of nanoluciferase (Nluc) co-transcribed with neomycin resistance (Neo^R^) driven by enolase promoter (15) (Figure 1C). Similar to previous work, the Nluc and Neo proteins were separated by a split peptide motif (P2A) (14). This tagging construct was co-transfected with a CRISPR/Cas9 plasmid containing a single guide RNA sequence (sgRNA) located in the C-terminus of the gene. To create parasites with INS1 fused to green fluorescent protein (GFP), we replaced the 3HA tag with GFP and used the same INS1 sgRNA plasmid (Figure 1D).

**Figure 1.**
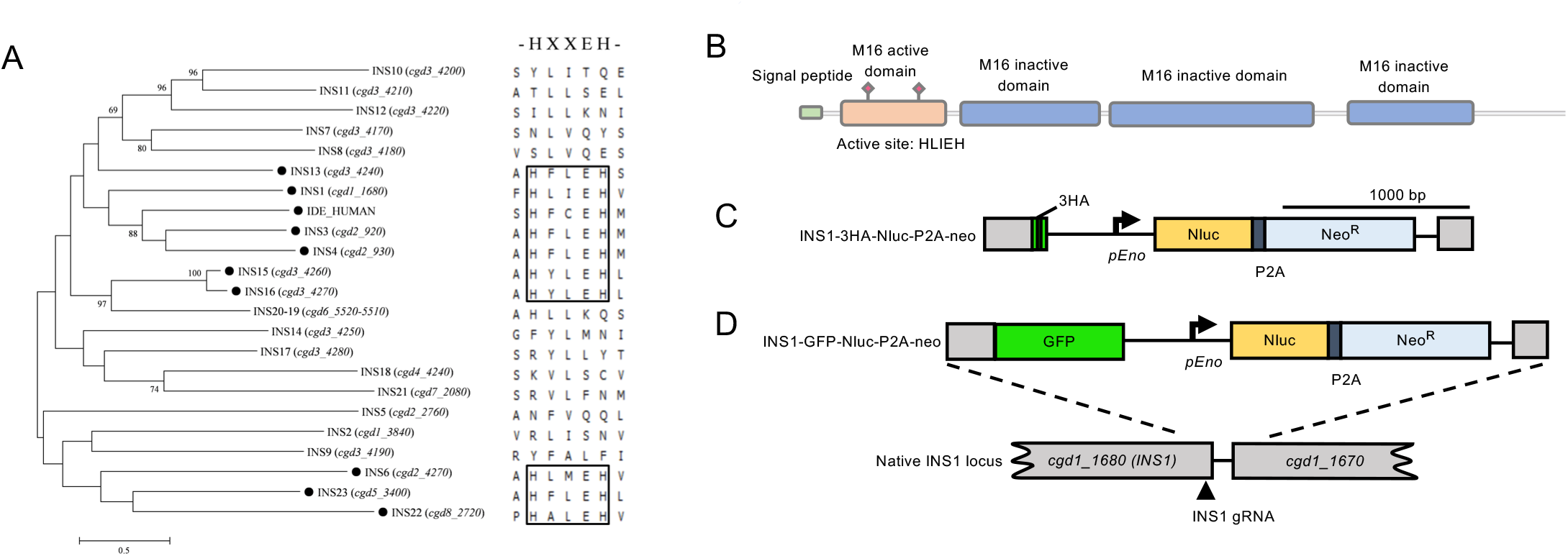
Insulinase-like proteases in *Cryptosporidium parvum* and construction of *C. parvum* INS1-3HA and INS1-GFP strains. (A) Phylogenetic relationship of *C. parvum* insulinase-like protease family and human insulinase. The tree was constructed by a maximum likelihood analysis with 1,000 replications for bootstrapping. The active site “HXXEH” in each INS was defined by multiple alignment. The black dot indicates proteins with a predicted active site. Scale = 50 changes per 100 residues. (B) Domain architecture of INS1 showing the presence of one M16 active domain containing an active site “HLIEH” and three inactive domains. (C) Diagram of INS1-3HA tagging strategy. Construct was designed to add a 3HA tag and Nluc-P2A-Neo^R^ cassette at the C-terminus of INS1 (*cgd1_1680*). P2A, split peptide; INS1 gRNA marks the site of guide RNA homology. (D) Diagram of INS1-GFP tagging strategy. Construct was designed to add a GFP tag and Nluc-P2A-Neo^R^ cassette at the C-terminus of INS1 (*cgd1_1680*). P2A, split peptide; INS1 gRNA site of guide RNA homology.

### Generation of stable transgenic INS1 parasites

Excysted sporozoites were electroporated with INS1-3HA-Nluc-P2A-neo tagging plasmid and CRISPR/Cas9 plasmid containing the INS1 sgRNA. Following electroporation, sporozoites were used to infect Ifngr1^-/-^ mice. Once oocyst shedding reached a peak level (13 dpi), and similar to mice infected with wild type oocysts, all of these mice succumbed to infection by day 20, indicating the tagged lines are not attenuated (Figure S1). Feces were collected from the first round of mice and a slurry was gavaged into a group of NSG mice to obtain the larger numbers of transgenic oocysts for purification. All mice were treated with 16 g/L paromomycin drinking water for selection of stable transgenic parasites (Figure 2A), as described previously (15). The signal of luminescence from the nLuc gene, increased in fecal pellets from 6 dpi (days post infection) with a peak value on 12 dpi (Figure 2C). Quantification of oocysts in feces by qPCR paralleled the increase in luciferase activity (Figure 2D). PCR analysis of oocysts collected from the mice confirmed that the INS1—3HA tagging cassette had correctly inserted into the *INS1* locus as shown using diagnostic primers that only amplify from the correct transgenic arrangement (Figure 2B and 2E). To obtain a tagged INS1-GFP strain, we used the same strategy to amplify the INS1-GFP transgenic parasites. In the first round of infection in Ifngr1-/- mice, all of the animals succumbed by day 20, indicating the tagged line was not attenuated (Figure S1). In the second round of amplifcation in NSG mice, luminescence and oocysts numbers measured from infected mice feces increased on 6 dpi and remained elevated one month (Figure 2G, 2H). PCR analysis confirmed that the INS1-GFP had correctly inserted into the *INS1* locus (Figure 2F and 2I). Fecal pellets were collected every day from 12-30 dpi for purification of transgenic parasites.

**Figure 2.**
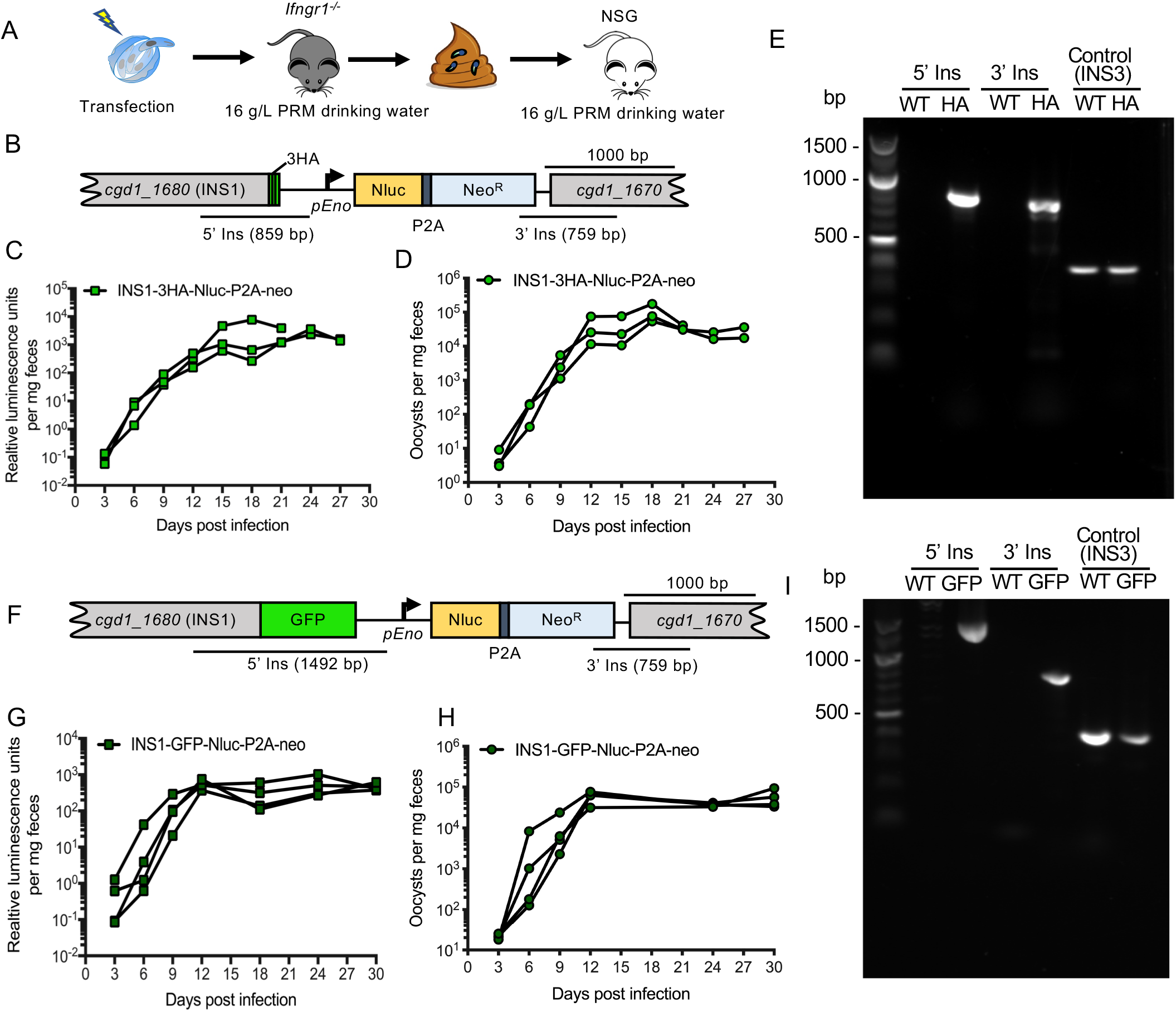
Amplification of transgenic parasites in immunocompromised mice. (A) Amplification strategy for obtaining tagged INS1 parasites. Approximately 5 × 10^7^ sporozoites were co-transfected with 50 μg tagging plasmid and 30 μg CRISPR/Cas9 plasmid per cuvette. Each Ifngr1^-/-^ mouse was gavaged with 200 μl 8% sodium bicarbonate solution 5 min before being infected by gavaged with 2.5 × 10^7^ transfected sporozoites in 200 μl of DPBS. A second round of selection was conducted in NSG mice. Each mouse was gavaged with fecal slurry containing 20,000 oocysts obtained at 13 dpi from the first round of selection. All mice received 16 g/L paromomycin drinking water from the first day post-infection (dpi) for the duration of the experiment. (B) Diagram of the INS1-3HA tagged locus in stable transgenic parasites. *C. parvum* were co-transfected with INS1-3HA-Nluc-P2A-neo tagging plasmid and CRISPR/Cas9 plasmid containing an INS1 sgRNA specific to the *INS1* locus. (C) Relative luminescence per mg of feces from transgenic *C. parvum* oocysts. Each data point represents a single pellet and each connecting line represents an individual infected NSG mouse from round two amplification of transfected parasites. (D) The number of oocysts per mg of feces was measured by qPCR. Each data point represents a single pellet and each connecting line represents an individual NSG mouse from round two amplification of transfected parasites. (E) PCR analysis of INS1-3HA oocysts amplified in mice from round two. WT, wild type. HA, INS1-3HA transgenic parasites. The product 5’ Ins is specific for the 5’ CRISPR targeting site of INS1-3HA. The product 3’ Ins is specific for the 3’ CRISPR targeting site of INS1-3HA. Control product is specific to the INS3 locus. Primers defined in Table S1. (F) Diagram of the INS1-GFP tagged locus in stable transgenic parasites. *C. parvum* were cotransfected with the INS1-GFP-Nluc-P2A-neo tagging plasmid and CRISPR/Cas9 plasmid containing an INS1 gRNA specific to the *INS1* locus. (G) Relative luminescence per mg of feces from transgenic *C. parvum* oocysts. Each data point represents a single pellet and each connecting line represents an individual NSG mouse infected with INS1-GFP parasites from round two. (H) The number of oocysts per mg of feces was measured by qPCR. Each data point represents a single pellet and each connecting line represents an individual NSG mouse infected with INS1-GFP parasites from round two. (I) PCR analysis of INS1-GFP oocysts amplified in mice from round two. WT, wild type. GFP, INS1-GFP transgenic parasites. The product 5’ Ins, is specific for the 5’ CRISPR targeting site of INS1-GFP. The product 3’ Ins, is specific for the 3’ CRISPR targeting site of INS1-GFP. Control, product is specific to the *INS3* locus. Primers defined in Table S1.

### INS1 is expressed in macrogamonts

In order to examine the expression of *INS1* during intracellular development of *C. parvum in vitro*, we infected HCT-8 cells with wild-type oocysts and tested transcription by reverse transcription-quantitative PCR (RT-qPCR) at different time points. We used two slightly different sets of RNA samples designed to cover the time range of the development of asexual and sexual stages, as described previous (8). The *INS1* gene showed no transcription before 30 hpi, but was upregulated at 36 hpi and reached the highest level at 48 hpi, before declining at 72 hpi (Figure 3A). The sexual stages of *C. parvum* first appear ~36 hpi (7, 8) and the positive signal for INS1 at this time point indicates that *INS1* was expressed in the either macrogamonts or microgamonts. To visualize the stage specific expression and localization of INS1, we performed immunofluorescence assays (IFA) using INS1-3HA and INS1-GFP transgenic parasites. Neither the anti-HA antibody nor anti-GFP antibody detected any staining in asexual stages of *C. parvum*, consistent with the absence of transcription during these stages. Instead, INS1 was only found in macrogamont stage where it appeared as a punctate staining, while microgamonts remained negative (Figure 3B, 3C).

**Figure 3.**
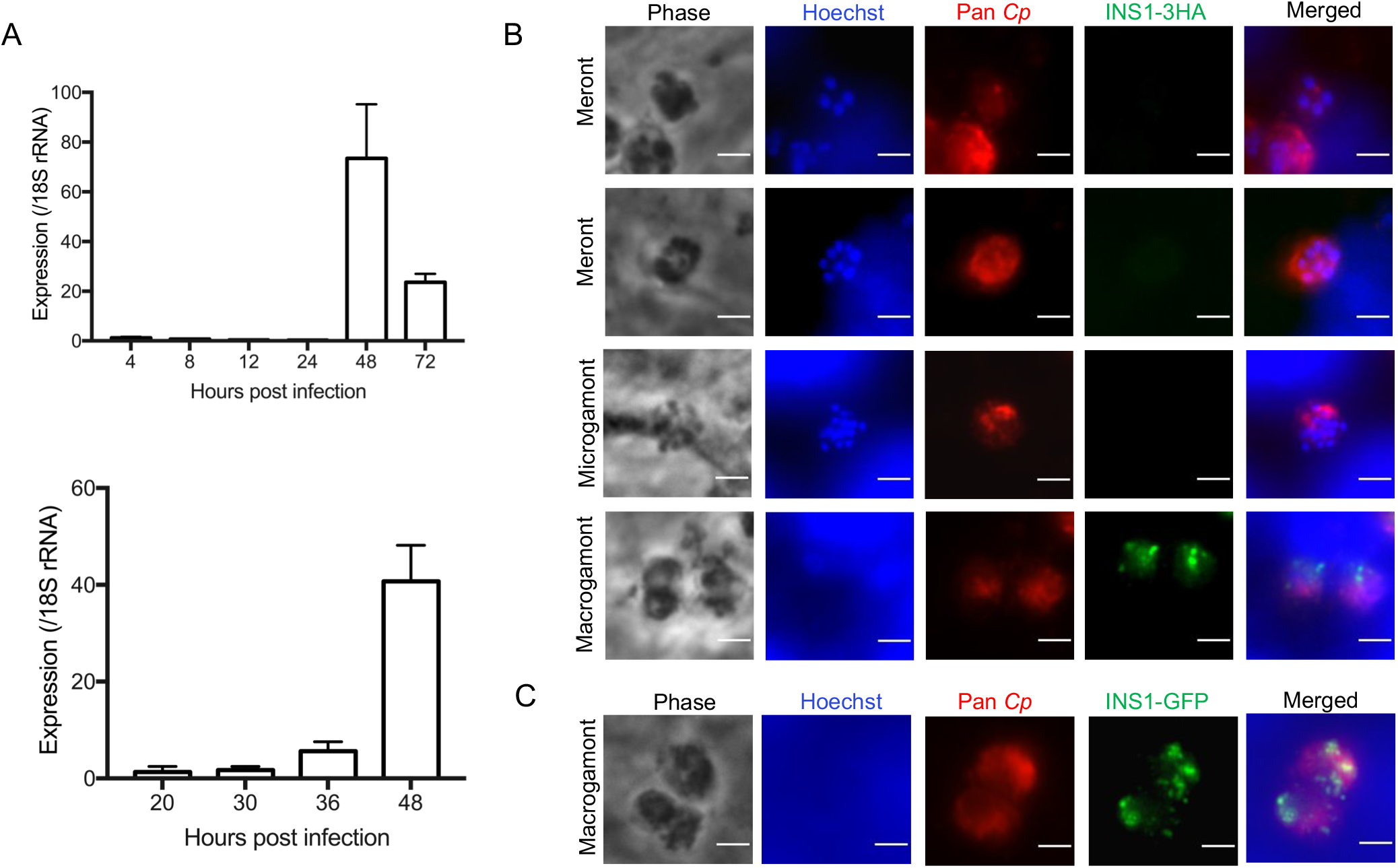
Transcription and expression of *C. parvum* INS1 *in vitro*. (A) Relative transcription level of the INS1 gene (*cgd1_1680*) at specified times post-infection as determined by reverse transcription-quantitative PCR. HCT8 cells were infected with *C. parvum* oocysts, cultured for specific time point, and RNA was collected from three wells per time point. Gene expression profiles are from two separate experiments with different time points. Data from the *Cryptosporidium* 18S rRNA gene were used in data normalization. Values are plotted as the means ± SD. (B) Immunofluorescence staining of transgenic INS1-3HA parasites. HCT-8 cells were infected with INS1-3HA oocysts. After 48 hpi, coverslips were fixed and stained with rat anti-HA followed by goat anti-rat IgG Alexa Fluor 488; rabbit Pan Cp followed by goat anti-rabbit IgG Alexa Fluor 568; and Hoechst for nuclear staining. Scale bars = 2 μm. (C) Immunofluorescence staining of INS1-GFP parasites. HCT-8 cells were infected with INS1-GFP oocysts. After 48 hpi, coverslips were fixed and stained with rabbit anti-GFP followed by goat anti-rabbit IgG Alexa Fluor 488; rat Pan Cp followed by goat anti-rat IgG Alexa Fluor 568; and Hoechst for nuclear staining. Scale bars = 2 μm.

Although there are a limited number of stage-specific proteins that have been characterized in *C. parvum*, we investigated two antibodies that have been previously shown to detect proteins that are expressed in macrogamonts. The punctate structures stained positively for INS1 did not colocalize with mAb 4D8, which recognizes a prominent filament structure of macrogamonts (11) (Figure 4A). We also examined costaining with mAb OW50 that stains cytoplasmic inclusions called wall forming bodies that are released to form the oocyst wall (30). INS1 was in distinct clusters from OW50; however, these were often in close proximity to each other (Figure 4B). To explore the ultrastructural localization of INS1, INS1-GFP transgenic parasites were examined the immunoelectron microscopy. INS1 was located in small vesicles within macrogamonts and these positive compartments were often in proximity with large electron dense vesicles (Figure 4C). Based on its timing of expression and localization, these findings suggest that INS1 might play some role in the development of macrogamonts.

**Figure 4.**
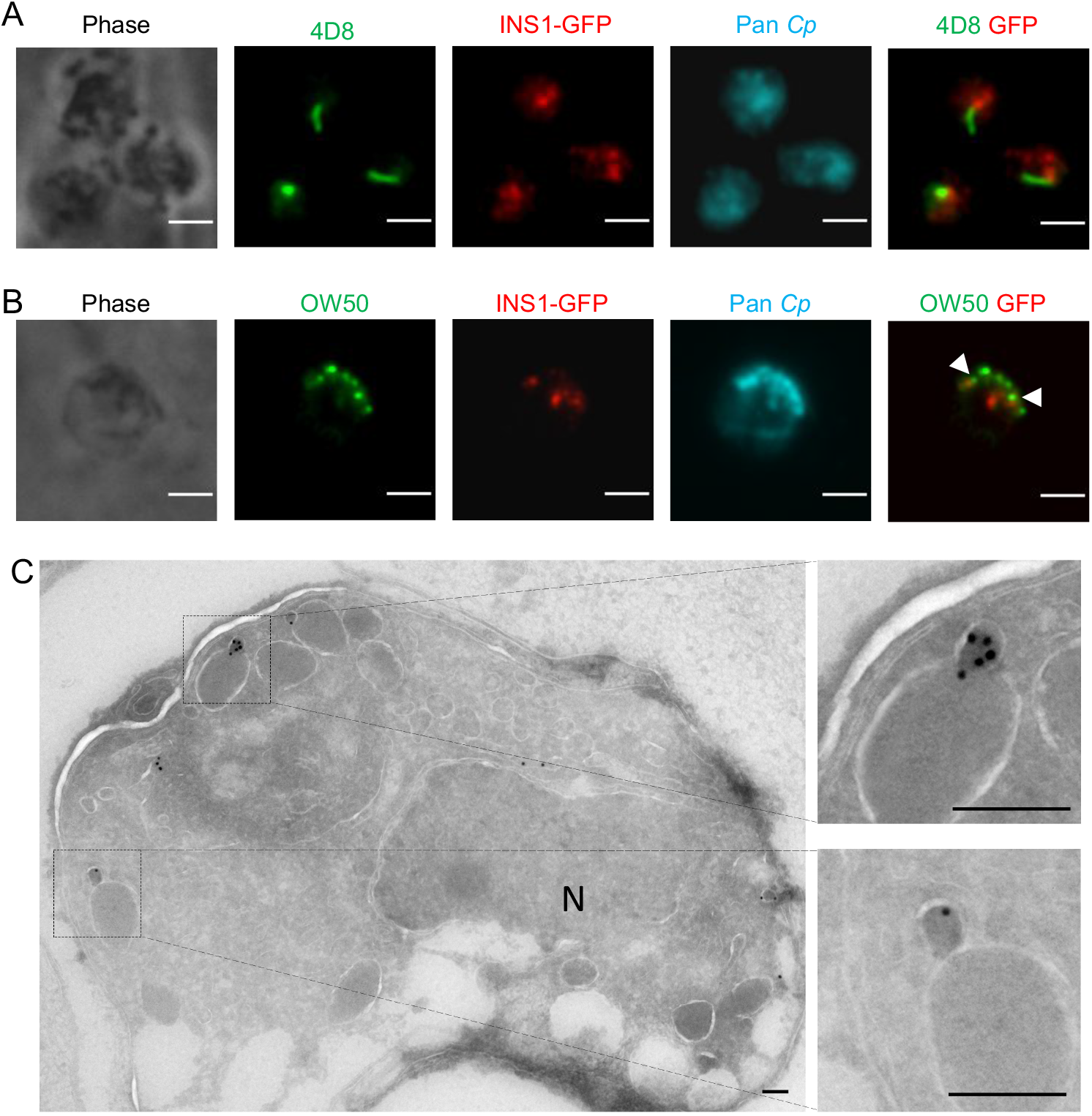
Expression of INS1 in different life cycle stages of *C. parvum*. (A) Immunofluorescence staining of macrogamont specific mAb 4D8 in INS1-GFP parasites. HCT-8 cells were infected with INS1-GFP oocysts. After 48 hpi, coverslips were fixed and stained with mouse mAb 4D8 followed by goat anti-mouse IgM Alexa Fluor 488; rabbit anti-GFP followed by goat anti-rabbit IgG Alexa Fluor 568; rat Pan Cp followed by goat anti-rat IgG Alexa Fluor 647; and Hoechst for nuclear staining. Scale bars = 2 μm. (B) Immunofluorescence staining of OW50 in INS1-GFP parasites. Arrow heads indicate staining of OW50 positive vesicles that were in close proximity with INS1. HCT-8 cells were infected with INS1-GFP oocysts. After 48 hpi, coverslips were fixed and stained with mouse mAb OW50 followed by goat anti-mouse IgG Alexa Fluor 488; rabbit anti-GFP followed by goat anti-rabbit IgG Alexa Fluor 568; rat Pan Cp followed by goat anti-rat IgG Alexa Fluor 647; and Hoechst for nuclear staining. Scale bars = 2 μm. (C) Transmission electron micrographs of macrogamont of INS1-GFP parasites. HCT-8 cells were infected with INS1-GFP oocysts. After 48 hpi, cells were fixed and stained with rabbit anti-GFP followed by 18 nm colloidal gold goat anti-rabbit IgG. Two images on the right are enlarged sections of the left image as indicated by the dotted lines. N, nucleus. Scale bars=200 nm.

### Decreased oocysts shedding in Δ*ins1* parasites

To investigate INS1 function in *C. parvum*, we generated *INS1* knockout (Δ*ins1*) parasites using CRISPR/Cas9. The *INS1* gene was replaced with a mCherry tag driven by the *C. parvum* actin promoter in a construct that also contained the Nluc-P2A-neo^R^ selection cassette described above (Figure 5A). To assure complete removal of the *INS1* gene, we used two sgRNA sequences to degenerate double stand DNA breaks that flank the gene (Figure 5A). Sporozoites were electroporated with the INS1-mCh-Nluc-P2A-neo-INS1 plasmid and a CRISPR/Cas9 plasmid containing the two *INS1* sgRNAs (Figure 5A). Transfected sporozoites were gavaged into a group of Ifngr1^-/-^ mice treated with 16 g/L PRM in the drinking water. Unlike the tagged lines described above, this lines was attenuated and none of the Ifngr1^-/-^ mice succumbed over a 30 day period (Figure S1). After 13 dpi, the feces containing the Δ*ins1* oocysts were collected and gavaged into a group of immunocompromised mice. For this second round infection, we were interested in confirming the attenuation of this line and so we used GKO mice, which like Ifngr1^-/-^ mice are highly sensitive to *C. parvum* infection (Figure 5C). PCR analysis confirmed that oocysts shed by GKO mice were Δ*ins1* parasites based on correct 5’ and 3’ flanking sequences created by loss of the *INS1* gene and insertion of the mCherry selection cassette in its place (Figure 5D). Additionally, primers to the open reading frame of INS1 failed to amplify a product from the knockout line although they easily detected the expected fragments from wild type parasites (Figure 5D).

**Figure 5.**
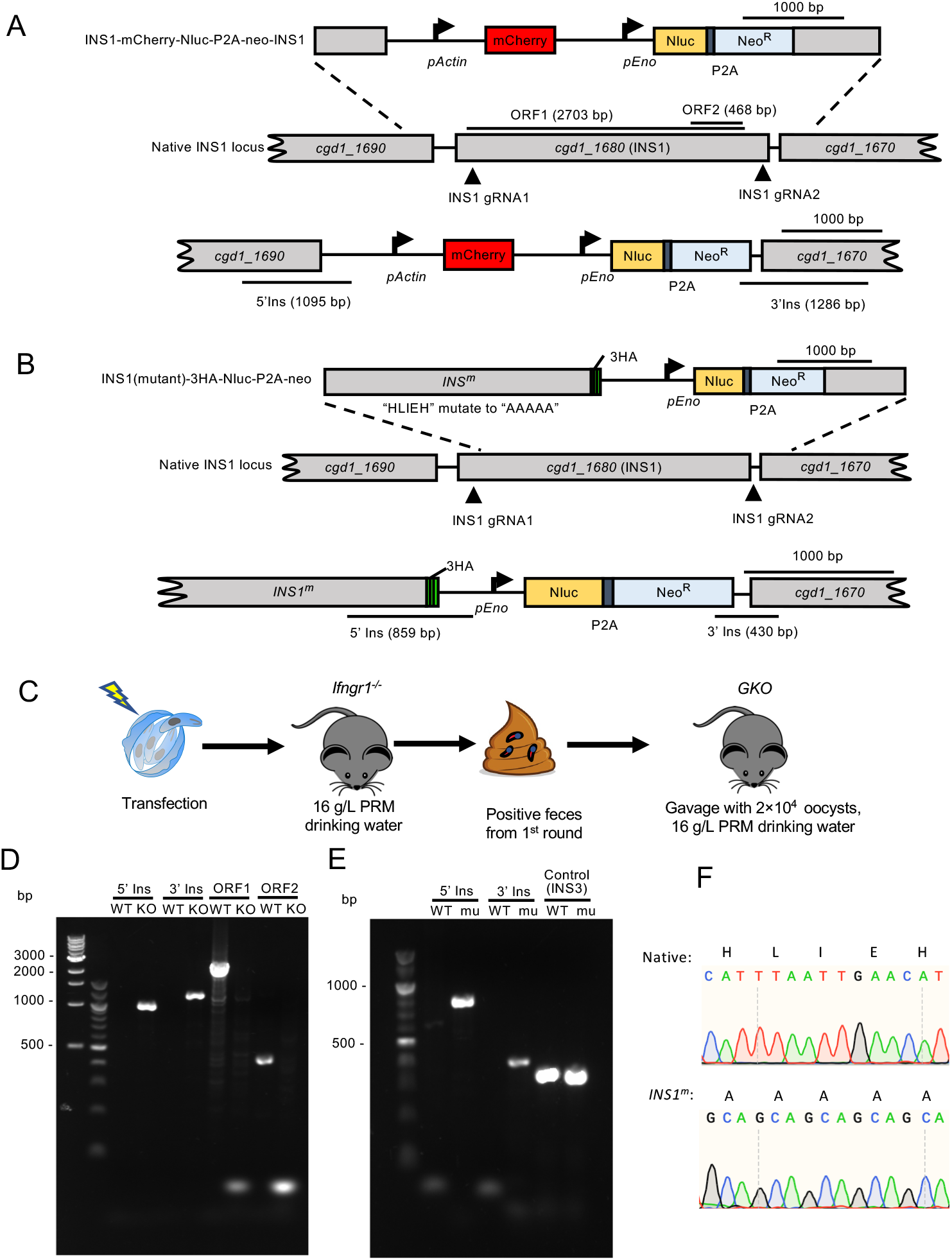
Selection of Δ*ins1* parasites and *INS1^m^* transgenic parasites in immunocompromised mice. (A) Diagram of the strategy to construct Δ*ins1* transgenic parasites by co-transfection with INS1-mCherry-Nluc-P2A-neo-INS1 targeting plasmid and CRISPR/Cas9 plasmid containing two INS1 sgRNA. The top line shows the targeting construct, the middle line the genomic locus, and the bottom line the successfully targeted transgenic locus. (B) Diagram of the strategy to construct *INS1^m^* active site mutants. Construct was designed to make an INS1 point mutation in which the active site “HLIEH” was mutated to “AAAAA” and add a 3HA tag and Nluc-P2A-Neo^R^ cassette at the C-terminus of INS1 (*cgd1_1680*). P2A, split peptide; INS1 gRNA site of guide RNA homology. (C) Selection strategy for obtaining Δ*ins1* or *INS1^m^* transgenic parasites. Transfected sporozoites were gavaged into Ifngr1^-/-^ mice treated with 16 g/L paromomycin in the drinking water. A second round of selection was conducted in GKO mice. Each mouse in round two was gavaged with fecal slurry containing 2 x 10^4^ oocysts collected at 18 dpi of first round of selection. (D) PCR analysis of Δ*ins1* oocysts obtained from the second round of amplification. WT, wild type. KO, Δ*ins1* parasite. The product 5’ Ins, is specific for the 5’ CRISPR targeting site of Δ*ins1* KO. The product 3’ Ins, is specific for the 3’ CRISPR targeting site of Δ*ins1*. The product ORF1 detects a 2,703 bp size fragment of the *INS1* open reading frame. The product ORF2, detects a 468 bp size fragment of the *INS1* open reading frame. (E) PCR analysis of Δ*ins1* oocysts obtained from the second round of amplification. WT, wild type. mu, *INS1^m^* parasite. The product 5’ Ins, is specific for the 5’ CRISPR targeting site of *INS1^m^*. The product 3’ Ins, is specific for the 3’ CRISPR targeting site of *INS1^m^*. Control, product is specific to the *INS3* locus. (F) Sequence electropherogram of PCR products from native INS1 (top) and active site mutant *INS1^m^* (bottom) transgenic parasites. Native, the amino acid and nucleotide sequence of active site in wilt type INS1 parasite. *INS1^m^*, the amino acid and nucleotide sequence of active site in *INS1^m^* parasite.

We then tracked the expansion of the Δ*ins1* parasites by luciferase measurements and oocyst shedding in the feces in the second round of mice. Both luminescence values and oocyst shedding increased on 6 dpi and reached to the peak at 12 dpi (Figure 6A). Moreover, mice infected by Δ*ins1* parasites continued to shed oocysts as long as one month, the longest time point we tracked (Figure 6B). In contrast, infection of GKO mice with similar numbers of wild type parasites led to illness after 12 dpi and none of the mice survived after 15 dpi (Figure 6B). The peak shedding with wild type oocysts reached 3.3 × 10^5^ oocysts per mg feces and this level was 5.7 times higher than shedding by Δ*ins1* mutants, a difference that was statistically significant.

**Figure 6.**
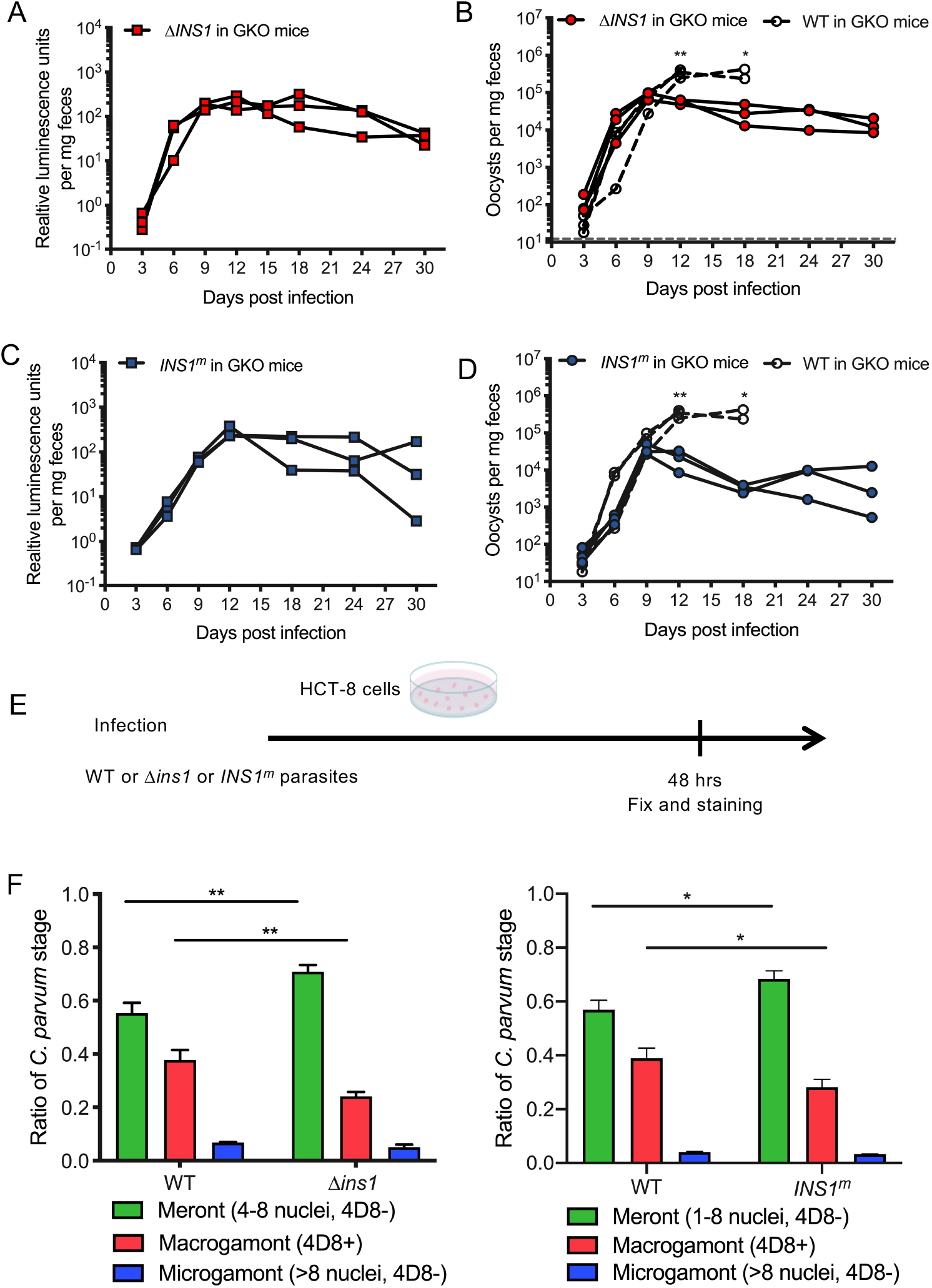
Influence of Δ*ins1* parasite and *INS1^m^* parasites on development of *C. parvum in vivo* and *in vitro*. (A) Relative luminescence of *C. parvum* per mg feces. Each red box represents a single pellet and each connecting line represents an individual GKO mouse infected with Δ*ins1* parasites from the second round of amplification. (B) The number of oocysts per mg of feces was measured by qPCR. Each red square represents a single pellet and each connecting line represents an individual GKO mouse infected with Δ*ins1* parasites from the second round of amplification. Each open box represents a single pellet and each connecting line represents an individual GKO mouse infected with 2 × 10^4^ WT oocysts. Statistical analysis was performed using unpaired Student’s t test for two sample comparison (*, *P* < 0.05; **, *P* < 0.01). (C) Relative luminescence of *C. parvum* per mg feces. Each blue box represents a single pellet and each connecting line represents an individual GKO mouse infected with INS1-mutant parasites from the second round of amplification. (D) The number of oocysts per mg of feces was measured by qPCR. Each blue square represents a single pellet and each connecting line represents an individual GKO mouse infected with INS1 mutant parasites from the second round of amplification. Each open box represents a single pellet and each connecting line represents an individual GKO mouse infected with 2 × 10^4^ WT oocysts. Statistical analysis was performed using unpaired Student’s t test for two sample comparison (*, *P* < 0.05; **, *P* < 0.01). (E) Outline of the experimental protocol to analyze growth of WT or Δ*ins1* or *INS1^m^* parasites in HCT8 cells. *C. parvum* WT or Δ*ins1* or *INS1^m^* parasites were used to infect HCT8 cells. After 48 hpi, wells were washed, fixed and labeled with different antibodies. For WT parasites, coverslips were stained with mouse mAb 4D8 that detects macrogamonts followed by goat anti-mouse IgM Alexa Fluor 488; rabbit Pan Cp followed by goat anti-rabbit IgG Alexa Fluor 568; and Hoechst for nuclear staining. For Δ*ins1* parasites, coverslips were stained with mouse 4D8 followed by goat anti-mouse IgM Alexa Fluor 488; rat anti-mCherry followed by goat anti-rat IgG Alexa Fluor 568; rabbit Pan Cp followed by goat anti-rabbit IgG Alexa Fluor 647; and Hoechst for nuclear staining. For *INS1^m^* parasites, coverslips were stained with mouse 4D8 followed by goat anti-mouse IgM Alexa Fluor 488; rabbit Pan Cp followed by goat anti-rabbit IgG Alexa Fluor 568; rat anti-HA followed by goat anti-rat IgG Alexa Fluor 647; and Hoechst for nuclear staining. (F) Quantification of life cycle stages of wild type, Δ*ins1* (left) or *INS1^m^* (right) parasites. Meronts were identified by their content of 4-8 nuclei; Macrogamont were identified by labeling with mAb 4D8; microgamonts were identified by many small nuclei (~16 nuclei). Each time point represents the average of three biological replicates. The number of parasites was counted from 50 fields of view with a 100 × oil objective. Values are plotted as the means ± SD. Statistical analysis was performed using unpaired t test for two samples comparison (*, *P* <0.05; **, *P* <0.01).

### INS1 requires active protease activity for function

We further tested whether INS1 requires its protease activity for function. INS1 contains the active site “HLIEH” that is a conserved motif in M16 metalloproteases. We used CRISPR/Cas9 genome editing to alter the INS1 active site from “HLIEH” to “AAAAA” while also adding a C-terminal 3HA tag (Figure 5B). Replacement of the endogenous locus with this active site mutant template (*INS1^m^*) was guided by two sgRNAs sequences located in the N-terminus and 3’UTR of the gene (Figure 5B). Sporozoites were electroporated with the INS1(mu)-3HA-Nluc-P2A-neo plasmid and a CRISPR/Cas9 plasmid containing two *INS1* gRNAs and oocysts were amplified as before. Similar to the Δ*ins1* knockout, this active site mutant line was attenuated and none of the Ifngr1^-/-^ mice succumbed over a 30 day period (Figure S1). To confirm this attenuation, we used GKO mice for the second round of infection. PCR analysis demonstrated that oocysts shed by GKO mice contained the correct 5’ and 3’ insertion of the repair template integrated into the *INS1* locus (Figure 5E). Sanger DNA sequencing of PCR products amplified from *INS1^m^* parasites confirmed replacement of the wild type copy with the mutated active site in oocysts recovered from the GKOP mice (Figure 5F). The *INS1^m^* active site mutants grew with slower kinetics than wild type parasites, similar to the Δ*ins1* parasites described above (Figure 6C). The *INS1^m^* active site mutant was also attenuated in mice and infection resulted in significantly lower shedding in infected GKO mice that survived for up to one month. Moreover, the peak shedding of wild type oocysts was 9.2 times higher than *INS1^m^* oocysts (Figure 6D).

### Inhibition of macrogamont maturation in Δ*ins1* and *INS1^m^* parasites *in vitro*

To test the growth abilities of the knockout and active site mutants *in vitro*, we infected HCT-8 cells with 10^4^ oocysts of each of the mutant *vs*. wild type parasites and returned them to culture to allow development. After 48 hpi, we fixed cells and labeled them with mAb 4D8 to detect macrogamonts as well as Pan Cp antibody to detect all stages. Life cycle stages were then defined based as follows: meronts were classified by four to eight nuclei but no 4D8 labeling, macrogamonts were classified by a single nucleus and were positive 4D8 labeling, and microgamonts were classified by having 16 nuclei but no 4D8 labeling. Wild type parasite cultures showed 55% parasites were meronts, 38% parasites were macrogamonts and 7% parasites were microgamonts at 48 hpi. In contrast, Δ*ins1* cultures showed significantly fewer macrogamonts (24%) with an increase in the proportion of meronts (71%) (Figure 6F). Similarly, HCT-8 cultures infected with the active site *INS1^m^* mutant parasites showed significantly fewer macrogamonts (28%) with an increase in the proportion of meronts (68%) (Figure 6F). Collectively, these findings indicate that INS1 functions to facilitate macrogamont development and in its absence oocyst formation is impaired, leading to attenuation *in vivo*.

## Discussion

The *C. parvum* genome contains an expanded family of 22 M16 metalloproteases, the majority of which have unknown functions. Here we focused on INS1, which is a clan M16A metalloprotease most similar to human insulinase. INS1 contains a signal peptide and an active domain containing the catalytic site HXXEH followed by several inactive domains. Unlike previously described INS proteins that are expressed early in development in *C. parvum*, INS1 was expressed exclusively in macrogamonts where it localized to small vesicular structures in the cytosol. Deletion of INS1, or replacement with an active site mutant, resulted in reduced formation of macrogametocytes *in vitro* and lower oocysts shedding *in vivo*. Our studies reveal that INS1 likely participates in macrogamont formation and in its absence the parasite forms fewer oocysts *in vivo* and is attenuated in immunocompromised mice.

Metalloproteases are widespread in biology and they have been classified into 16 clans that are summarized in the MEROPS database (https://www.ebi.ac.uk/merops/). M16 metalloproteases are characterized by the presence of a zinc binding motif consisting of the sequence HXXEH (22). M16A and M16C family member contain four domains, only one of which contains the active catalytic site (22). The best known M16 protease is human insulinase that cleaves variety type of small peptides in different type of cells, consistent with multiple roles of this enzyme (31). Apicomplexan parasites contain numerous M16 metalloproteases, although their roles have only partially been investigated. Two M16C proteases have been described in *Plasmodium falciparum:* falcilysin is involved in hemoglobin degradation and processing of the apicoplast import transit peptides (26, 32), an activity that is also catalyzed by *PfSPP* (33). *Toxoplasma gondii* contains 11 M16 orthologues and several of M16A enzymes found in secretory compartments have been previously studied (22). Toxolysin 1 is found in rhoptries (23), while Toxolysin 4 is found in micronemes (24). Both proteases are themselves processed during maturation, but their substrates and roles these proteases play in their respective secretory compartments are undefined. Toxolysin 4 is refractory to gene disruption (24), while loss of toxolysin 1 has no effect on growth *in vitro* or virulence *in vivo* (23). Toxolysin 3 that is encoded by TGME49_257010 is most similar to INS1 by sequence homology. Toxolysin 3 is highly expressed in oocysts and is annotated as a “sporozoites development protein” (https://toxodb.org/toxo/app). In contrast, the INS1 is expressed exclusively in macrogamonts, indicating that these M16 metalloproteases have different biological roles.

Of the 22 M16 metalloproteases in encoded the *C. parvum* genome, 18 of them belong to the M16A family and several of these have previously been studied including INS5, INS15, and INS20-19. INS15 and INS20-19 contain signal peptides suggesting they are in the secretory pathway, while INS5 does not have a predicted signal peptide (27–29). INS20-19 is expressed in sporozoites and localized to an apical compartment (27), while INS15 is expressed in a middle-anterior compartment in sporozoites and merozoites (28). In contrast, INS5 is expressed at lower levels in sporozoites and increases to peak at 36-48 hpi (29). INS5 is present in a punctate pattern in sporozoites and during merogony (29). Antibodies to these proteins have been shown to inhibit parasite invasion *in vitro* (27–29), suggesting they play some role in processing or maturation of substrates involved in host cell recognition. INS1 is a classic M16A protease containing a signal peptide and an active motif “HLIEH” in the N-terminal domain followed by three domains that lack this catalytic motif. It also contains a signal peptide, but is not expressed in sporozoites nor merozoites, and hence is unlikely to function in host cell interactions. INS1 is encoded by the *cgd1_1680* gene in *C. parvum* and orthologues to this gene are present in other *Cryptosporidium* spp. (34), suggesting it has a conserved function.

INS1 was not detected in sporozoites or during merogony but rather expression was strongly upregulated at 36 to 48 hpi when the sexual stages are formed (12). Consistent with this pattern, previous studies have shown that INS1 and INS3 (encoded by *cgd2_920*) are highly expressed in macrogamonts *in vivo* and *in vitro* culture (7). Tagging with the HA epitope or a GFP fusion indicated that INS1 is expressed exclusively in macrogamonts where it is localized to small vesicular structures in the cytosol. INS1 was not colocalized with 4D8 antibody that recognizes a unique striated fiber in macrogamonts (11). Additionally, INS1 only partially colocalized with OW50 that recognizes large punctate vesicles in the cytosol of macrogamonts and the wall in mature oocysts (35). When examined by immunoelectron microscopy, INS1 was found in small vesicular structures that were often adjacent to large electron-dense vesicles. These large vesicles resemble wall forming bodies that have reported that contain proteins involved in formation of outer oocysts layer (36). Attempts to label OW50 in combination with INS1 by immunoelectron microscopy were not successful. Nonetheless, the proximity of small vesicles containing INS1 with the larger wall forming bodies suggests that INS1 may process some component of the oocyst wall during maturation in the secretory system. Like other insulinase enzymes (21), INS1 is predicted to have a small folded barrel containing the active site that is only capable of accommodating small peptides ranging from 3-6 kDa. As such INS1 is likely to process small peptides, possibly resulting from trimming of secretory proteins by other peptidases in the secretory pathway. Confirmation of such a role awaits identification of potential substrates of INS1.

Deletion of INS1, or replacement with the *INS1^m^* active site mutant, did not affect asexual replication *in vitro*, nor completely block development, indicating that INS1 is not essential for growth. Instead, loss of INS1 resulted in reduced formation of macrogamonts *in vitro* and lower oocysts shedding *in vivo*. In human insulinase, replacement of E^111^ in the active site “HXXEH” with Q^111^ inactivates catalysis (37, 38). Similarly, the insulinase homologue Iph1 in yeast loses catalytic activity when the active site E^71^ is changed to D^71^ (39). Although we have not formally demonstrated that the INS1^m^ enzyme has lost catalytic activity, the complete replacement of the “HXXEH” site with “AAAAA” is consistent with that interpretation. Hence, the similar phenotypes of the Δ*ins1* strain and the *INS1^m^* active site mutants suggests that the observed phenotype of the knockout is due to loss of catalytic activity. We also observed that the defect in the *INS1^m^* active site was approximate 2 times more severe in terms of reduced oocyst shedding than the complete deletion, suggest the expression of an inactive enzyme has a dominant negative effect that is greater than the loss of the enzyme. Both the Δ*ins1* knockout out and the INS1^m^ active site mutant were less virulent in GKO mice and reduced oocyst shedding was associated with survival beyond 12 days with mice infected with wild type parasites succumb to infection. The lower virulence of the INS1 mutants is likely an indirect consequence of reduced macrogamonts, fewer oocysts and less reinfection, since the pathology of infection is normally associated with multiple rounds of merogony.

Our studies reveal that INS1 participates in some aspect of macrogamont formation or maturation and that its catalytic function is necessary for optimal formation of oocysts. *Cryptosporidium* is transmitted through fecal-oral route by ingestion viable oocysts in contaminated water or food (40). Normally low doses of oocysts can cause diarrhea in humans or animals and the infected host can continue to shed a large quantity of oocysts over a period of several weeks (41). Infections are more severe in young children, where cryptosporidiosis is a major cause of debilitating diarrheal disease (42). The absence of effective treatments or vaccination makes it difficult to manage infections in these susceptible populations. Here we observed that loss of INS1 in results in attenuated in oocyst shedding and decreased virulence in immunocompromised mice. In addition to providing insight into the development of sexual stages and oocyst formation, such mutants could provide an approach for vaccination by challenge with attenuated strains designed to induce immunity without causing disease.

## Methods

### Animal studies

Animal studies on mice were approved by the Institutional Animal Studies Committee (School of Medicine, Washington University in St. Louis). The Ifngr1^-/-^ mice (Jackson Laboratories #003288), Ifng^-/-^ mice (referred to as GKO) (Jackson Laboratories #002287) and *Nod scid gamma* mice (referred to as NSG) (Jackson Laboratories #005557) were bred in house at Washington University School of Medicine and were separated by sex after weaning. Ifngr1^-/-^ and GKO mice raised in our facility are highly susceptible to the strain of *C. parvum* used here and they shed high numbers of oocysts and routinely die within 10-12 dpi. As such, they are used to amplify transgenic strains following initial transfection of sporozoites. In contrast, NSG mice are more resistant and can be used for prolonger oocyst shedding, which occurs are lower peak levels. Mice were reared in a specific-pathogen-free facility on a 12:12 light-dark cycle and water ad libitum. For selection and amplification of transgenic *C. parvum* parasites, 8-12 weeks old mice were used and water was replaced with filtered tap water containing 16 g/L paromomycin sulfate salt (Sigma). During the course of infection, animals that lost more than 20% of their body weight or became non-ambulatory were humanely euthanized.

### Phylogenetic analysis

The amino acid sequences of INS in *Cryptosporidium parvum* were extracted from CryptoDB (https://cryptodb.org/cryptodb/app) and the human insulinase was extracted from Uniprot (https://www.uniprot.org). The phylogenetic tree was constructed based on these sequences. MUSCLE was used to align the concatenated sequences. Phylogenetic trees based on maximum likelihood were constructed with 1,000 replications for bootstrapping.

### Primers

All primers were synthesized by Integrated DNA Technologies and are listed in Table S1.

### Oocysts preparation and excystation

*Cryptosporidium parvum* oocysts were obtained from the Witola lab (University of Illinois at Urbana-Champaign). The *C. parvum* isolate (AUCP-1) was maintained by repeated passage in male Holstein calves and purified from fecal material, as described previously (43). Purified oocysts were stored at 4°C in 50 mM Tris-10 mM EDTA (pH 7.2) for up to six months before use. Before infection, 1×10^8^ purified oocysts were diluted into 1 ml of Dulbecco’s phosphate-buffered saline (DPBS; Corning Cellgro) and treated with 3 ml of 40% bleach (containing 8.25% sodium hypochlorite) for 10 min on ice. Oocysts were then washed 4 times in DPBS containing 1% (wt/vol) bovine serum albumin (BSA; Sigma) and resuspended in 1 ml DPBS with 1% BSA. For some experiments, oocysts were excysted prior to infection by incubating the oocysts with 0.75% (wt/vol) sodium taurocholate (Sigma) at 37°C for 60 min.

### HCT8 cell culture

Human ileocecal adenocarcinoma cells (HCT-8 cells; ATCC CCL-244) were cultured in RPMI 1640 medium (Gibco, ATCC modification) supplemented with 10% fetal bovine serum. The HCT-8 cells were determined to be mycoplasma-negative using the e-Myco plus kit (Intron Biotechnology).

### Gene expression analysis

HCT-8 cells were grown on 6-well culture plates and incubated 24 h before infection. Monolayers were infected with excysted sporozoites, washed twice with DPBS at 2 hpi, and fresh HCT-8 medium then was added. RNA was collected from three wells per time point in RLT buffer (Qiagen) plus 1% β-mercaptoethanol, homogenized using a QIAshredder column (Qiagen), and then stored at −80°C until further processing. RNA was extracted using the RNeasy minikit (Qiagen), treated with the DNA-free DNA removal kit (Thermo Fisher Scientific), and converted to cDNA using the SuperScript VILO cDNA synthesis kit (Thermo Fisher Scientific). Reverse transcription quantitative PCR (RT-qPCR) was performed using a QuantStudio 3 realtime PCR system (Thermo Fisher Scientific) with SYBR Green JumpStart Taq ReadyMix (Sigma) using primers in Table S1. The following conditions were used for RT-qPCR: priming at 95°C for 2 min followed by 40 cycles of denaturing at 95°C for 10 sec, annealing at 60°C for 20 sec, and extension at 72°C for 30 sec, followed by a melt curve analysis to detect non-specific amplification. Relative gene expression was calculated with the ΔΔ*C_T_* method (44) using *C. parvum* 18S rRNA as the reference gene.

### Gene tagging using CRISPR/Cas9

To provide a single guide RNA (sgRNA) plasmid for INS1, the plasmid pACT1:Cas9-GFP, U6:sgINS1 was generated by replacing the TK sgRNA in pACT1:Cas9-GFP, U6:sgTK plasmid (14) with a sgRNA which located in the C-terminus of the targeting the *C. parvum* INS1 gene (*cgd1_1680*), 158 bp before the stop codon, using Q5 site-directed mutagenesis (New England Biosciences). The INS1 sgRNA was designed using the Eukaryotic Pathogen CRISPR guide RNA/DNA Design tool (http://grna.ctegd.uga.edu). To generate a tagging plasmid, a portion of the INS1 C-terminus (253 bp) with the mutant protospacer adjacent motif (PAM) and INS1 3’UTR (117 bp) was amplified from *C. parvum* genome DNA. The triple hemagglutinin (3HA) epitope tag was amplified from pTUB1:YFP-mAID-3HA, DHFR-TS:HXGPRT (45). The previously described Nluc-P2A-neo^R^ reporter including the pUC19 backbone was amplified from TK-GFP-Nluc-P2A-neo-TK plasmid (14). The tagging plasmid pINS1-3HA-Nluc-P2A-neo was then generated by Gibson assembly of the components listed above. To generate a C-terminal tag with a green fluorescent protein (GFP) tag, the plasmid pINS1-GFP-Nluc-P2A-neo was constructed by swapping the 3HA with GFP from TK-GFP-Nluc-P2A-neo-TK plasmid using Gibson assembly of PCR-amplified fragments.

### Gene deletion using CRISPR/Cas9

To provide a plasmid for gene deletion, a second sgRNA U6:sgINS1(2) was inserted in the plasmid pACT1:Cas9-GFP, U6:sgINS1 to generate the plasmid pACT1:Cas9-GFP, dual, U6:sgINS1-KO. This plasmid contains two sgRNAs that flank the gene, one located at the C-terminus of INS1, 158 bp before the stop codon, and one is located at N-terminus of INS1, 55 bp after promoter. The targeting plasmid pINS1-mCh-Nluc-P2A-neo-INS1 was made by replacing the UPRT homologous flanks with INS1 homologous flanks (770 bp upstream and 852 bp downstream of *cgd1_1680*) from UPRT-mCh-Nluc-P2A-neo-UPRT (14) using Gibson assembly of PCR-amplified fragments.

### Generating point mutations using CRISPR/Cas9

To generate an active site mutant of INS1, the plasmid pACT1:Cas9-GFP, dual, U6:sgINS1-mu was generated by inserting two new sgRNA in pACT1:Cas9-GFP, U6:sgTK plasmid. This plasmid contains two sgRNAs that flank the gene, one located at the N-terminus of INS1, 5 bp after active site, and one is located at 3’UTR of INS1, 3 bp after stop codon. To generate the targeting plasmid, a portion of the C-terminus INS1 homologous flanks with flanking regions before active site (3081 bp of *cgd1_1680*) and 117 bp INS1 3’UTR was amplified from *C. parvum* genome DNA, PAM sequences were mutated to prevent re-cutting the repair DNA. The active site in INS1 “HLIEH” was then mutated to “AAAAA” using Q5 site-directed mutagenesis. The 3HA-Nluc-P2A-neo^R^ reporter including the pUC19 backbone was amplified from INS1-3HA-Nluc-P2A-neo plasmid. The targeting plasmid pINS1(mu)-3HA-Nluc-P2A-neo was generated by Gibson assembly of the components listed above.

### Transfection of *C. parvum* sporozoites

Oocysts (1.25 x 10^7^ per transfection) were excysted as described above, sporozoites were pelleted by centrifugation and resuspended in SF buffer (Lonza) containing 50 μg of tagging or targeting plasmids and 30 μg CRISPR/Cas9 plasmid in a total volume of 100 μl. The mixtures were then transferred to a 100 μl cuvette (Lonza) and electroporated on an AMAXA 4D-Nucleofector System (Lonza) using program EH100. Electroporated sporozoites were transferred to cold DPBS and kept on ice before infecting mice.

### Selection and amplification of transgenic parasites in immunodeficient mice

Three Infgr1^-/-^ mice was used for the first round of transgenic parasite selection. Each mouse was orally gavaged with 200 μl of saturated sodium bicarbonate, 5 min prior to infection. Each mouse was then gavaged with 2.5 x 10^7^ electroporated sporozoites. All mice received drinking water with 16 g/L paromomycin continuously from the first day post-infection (dpi), based on previously published protocols (15). Fecal pellets were collected begin at 9 - 15 dpi, after which animals were euthanized by CO_2_ asphyxiation according to the animal protocol guidelines. Fecal pellets were stored at −80°C for qPCR or at 4°C for luciferase assays or for isolating oocysts for subsequent infections.

A second round of amplification was performed by orally gavaging 3 - 4 NSG mice (used for isolating tagged strains) or GKO mice (used for amplifying knockout or mutation lines) using a fecal slurry from round one mice described above. Fecal slurries were produced by grinding pellets (from 13 dpi) in DPBS, then centrifuging at 200 rpm for 10 min to pellet large particulates. The supernatant was then diluted in DPBS to achieve a concentration of ~ 2 x 10^4^ oocysts in 200 μl DPBS then gavaged into one mouse. Similar to round one, the mice infected in round two were treated with 16 g/L paromomycin drinking water for the entirety of the experiment. Fecal pellets for qPCR and luciferase assay were collected every 3 days starting 12 dpi and stored at 4°C. For purification, fecal samples from all mice were pooled and oocysts extracted as previously described (46). Purified oocysts were stored in PBS at 4°C and used within six months of extraction.

### Luciferase assay

Luciferase assays were performed with the Nano-Glo Luciferase Assay kit (Promega). Mouse fecal pellets were collected in 1.7 ml microcentrifuge tubes, ground with a pestle, then 3-mm glass beads (Fisher Scientific) and 1 ml fecal lysis buffer (50 mM Tris pH7.6, 2 mM DTT, 2 mM EDTA pH 8.0, 10% glycerol, 1% Triton X-100 prepared in water) (47) were added to the tube. Tubes were incubated at 4°C for 30 mins, vortexed for 1 min, then spun at 16,000 × g for 1 min to pellet debris. The 100 μl supernatant was added split between two wells of a 96-well white plate (Costar 3610). Then, 100 μl of a 25:1 Nano-Glo Luciferase buffer to Nano-Glo Luciferase substrate mix was added to each well and the plate was incubated for 3 min at room temperature. Luminescence values were read on a Cytation 3 Cell Imaging Multi-Mode Reader (BioTek).

### Fecal DNA extraction and quantification of oocysts

DNA was extracted from fecal pellets using the QIAamp PowerFecal DNA kit (Qiagen) according to manufacturer’s protocols. Oocyst numbers were quantified using qPCR with the *C. parvum* GAPDH primers (Table S1), as described previously (14). A standard curve for *C. parvum* genomic DNA was generated by purifying DNA from a known number of oocysts and creating a dilution series. Reactions were performed on a QuantStudio 3 Real-Time PCR System (Thermo Fisher) with the amplification conditions as previously described (14).

### PCR identification of transgenic parasites

To check for the successful insertion of the target sequence into the *INS1* locus, PCR was performed on 1 μl purified fecal DNA using Q5 Hot Start High-Fidelity 2 × master mix (New England Biosciences) with primers listed in Table S1 at a final concentration of 500 nM each. PCR reactions were performed on a Veriti 96-well Thermal Cycler (Applied Biosystems) with the following cycling conditions: 98°C for 30 secs, followed by 35 cycles of 98°C for 15 secs, 60°C for 30 secs, and 72 °C for 2 mins, with a final extension of 72 °C for 2 mins. PCR products were resolved on 1.0% agarose gel containing GelRed (Biotium, diluted 1:10,000) and imaged on a ChemiDoc MP Imaging System (Bio-Rad).

### Indirect immunofluorescence microscopy

HCT-8 cells grown on coverslips were infected 24 h post-seeding with 1 × 10^5^ oocysts per well, then fixed with 4% formaldehyde at specific timepoints. The fixed samples were washed twice with DPBS and then permeabilized and blocked with DPBS containing 1% BSA and 0.1% Triton X-100 (Sigma). Primary antibodies were diluted in blocking buffer for staining: rat anti-HA was used at 1:500, rabbit anti-GFP was used at 1:1000, mAb 4D8 (hybridoma supernatant) was used at 1:20, mAb OW50 (11) was used at 1:10 and pan Cp (rabbit pAb or rat pAb) was used at 1:10,000. Cells were incubated with primary antibodies for 60 min at room temperature, washed three times with PBS, then incubated for 60 min at room in secondary antibodies conjugated to Alexa Fluor dyes (Thermo Fisher Scientific) diluted 1:1,000 in blocking buffer. Nuclear DNA was stained with Hoechst (Thermo Fisher Scientific) diluted 1:1,000 in blocking buffer for 15 min at room temperature, then mounted with Prolong Diamond Antifade Mountant (Thermo Fisher Scientific). Imaging was performed on a Zeiss Axioskop Mot Plus fluorescence microscope equipped with a 100×, 1.4 N.A. Zeiss Plan Apochromat oil objective lense and an AxioCam MRm monochrome digital camera. Images were acquired using Axiovision software (Carl Zeiss Inc.) and manipulated in ImageJ or Photoshop.

### Transmission Electron Microscopy

HCT-8 cells were infected with INS1-GFP parasites for 48 h, infected cells then were fixed in freshly prepared mixture of 4% paraformaldehyde and 0.05% glutaraldehyde (Polysciences Inc., Warrington, PA) in 100 mM PIPES/0.5 mM MgCl_2_ buffer (pH 7.2) for 60 min at 4°C. Samples were then embedded in 10% gelatin and infiltrated overnight with 2.3 M sucrose/20% polyvinyl pyrrolidone in PIPES/MgCl_2_ at 4°C. Samples were trimmed, frozen in liquid nitrogen, and sectioned with a Leica Ultracut UCT7 cryo-ultramicrotome (Leica Microsystems Inc., Bannockburn, IL). Ultrathin sections of 50 nm were blocked with 5% fetal bovine serum/5% normal goat serum for 30 min and subsequently incubated with rabbit anti-GFP antibody (Life Technologies Corp., Eugene, OR) for 60 min at room temperature. Following washes in block buffer, sections were incubated with goat anti-rabbit IgG (H+L) conjugated to 18 nm colloidal gold (Jackson ImmunoResearch Laboratories, Inc., West Grove, PA) for 60 min. Sections were stained with 0.3% uranyl acetate/2% methyl cellulose and viewed on a JEOL 1200 EX transmission electron microscope (JEOL USA Inc., Peabody, MA) equipped with an AMT 8 megapixel digital camera and AMT Image Capture Engine V602 software (Advanced Microscopy Techniques, Woburn, MA). All labeling experiments were conducted in parallel with controls omitting the primary antibody. These controls were consistently negative at the concentration of colloidal gold conjugated secondary antibodies used in these studies.

### Quantification and statistical analysis

All statistical analyses were performed in GraphPad Prism 8 (GraphPad Software) unless otherwise specified. When comparing the means of two groups at the same time point, we used an unpaired t test. For statistical analysis *P* values ≤ 0.05 were considered significant.

## Data Availability

All of the data are found in the manuscript or Supplemental Materials.

## Acknowledgements

We thank members of the Sibley laboratory for helpful comments Lisa Funkhouser-Jones for providing cDNA samples, and Soumya Ravindran for technical assistance, and Wandy Beatty, Microbiology Imaging Facility at Washington University in St. Louis, for performing the electron microscopy studies. This work was partially supported by a grant from the China Scholarship Council to RX and a grant from the National Institutes of Health (AI149309) to LDS.

## Supplemental Material

Table S1 – Oligonucleotides used in this study.

Figure S1. Survival curve of Ifngr1^-/-^ mice infected with different transgenic parasites.

For INS1-3HA, INS1-GFP, Δ*ins1* and *INS1^m^*, each Ifngr1^-/-^ mouse was gavaged with 2.5 × 10^7^ transfected sporozoites, and mice received 16 g/L paromomycin drinking water from the first day post-infection (dpi) for the duration of the experiment. In INS1-3HA and INS1-GFP group, three mice were gavaged, although only two of them became infected and the data plotted here are the composite survival of those two animals. In the Δ*ins1* and *INS1^m^* groups, three mice were gavaged, all of them were infected and the data shown are based on all three animals. For WT group, each Ifngr1^-/-^ mouse was gavaged with 2 × 10^4^ oocysts and four mice were used in the experiment.

